# Syntactic Sugars: Crafting a Regular Expression Framework for Glycan Structures

**DOI:** 10.1101/2024.02.01.578383

**Authors:** Alexander R. Bennett, Daniel Bojar

## Abstract

**Summary:** Structural analysis of glycans pose significant challenges in glycobiology due to their complex sequences. Research questions such as analyzing the sequence content of the α1-6 branch in *N*- glycans, are biologically meaningful yet can be hard to automate. Here, we introduce a regular expression system, designed for glycans, feature-complete, and closely aligned with regular expression formatting. We use this to annotate glycan motifs of arbitrary complexity, perform differential expression analysis on designated sequence stretches, or elucidate branch-specific binding specificities of lectins in an automated manner. We are confident that glycan regular expressions will empower computational analyses of these sequences.

**Availability and implementation:** Our regular expression framework for glycans is implemented in Python and is incorporated into the open-source glycowork package (version 1.1+). Code and documentation are available at https://github.com/BojarLab/glycowork/blob/master/glycowork/motif/regex.py.

## Main

Glycans are complex carbohydrates, key in many biological processes (Varki, 2017). Monosaccharides, linked together in specific patterns, are attached to proteins and lipids, influencing their function and properties. Diversity in glycans arises from different monosaccharide monomers and the ways in which these can be linked, leading to a vast array of possible structures even with a limited set of building blocks (Bojar *et al*., 2021).

The biological importance of glycans lies in motifs, distinct sequence patterns within the larger structure. For example, the sialyl Lewis^x^ motif, Neu5Acα2-3Galβ1-4(Fucα1-3)GlcNAc, is crucial for cell adhesion and is implicated in cancer metastasis (Jin and Wang, 2020), while high-mannose motifs are recognized by proteins from the immune system (Dommett *et al*., 2006), and blood group antigens, such as the ABO blood groups (Stanley *et al*., 2022), are determined by specific glycan motifs on red blood cells.

Understanding glycans, and their motifs, is vital for unraveling their roles in health and disease (Reily *et al*., 2019), and in developing targeted therapeutic strategies. While motifs drive biological functions, their position within the glycan is crucial for functionality. The simplest example of this is distinguishing terminal and internal forms of the same motif, which already affects biological properties, as many proteins or lectins only recognize one form (Bojar *et al*., 2022). More distal context effects, such as whether a motif is presented on the α1-3 or the, often not outstretched (Fogarty *et al*., 2020), α1-6 branch in *N*-glycans, also can have drastic effects, as most lectins have an arm preference for their binding motif (Li *et al*., 2020). Examples include MAL-I preferring Neu5Acα2-3 on the α1-6 branch, whereas SNA strongly favored Neu5Acα2-6 on the α1-3 branch. Especially this latter category of motifs, for which the larger sequence context is key, are hard to capture with existing methods.

Here, we present a regular expression (RegEx) system for glycans to search for, and extract, sequence patterns of arbitrary complexity. Regular expressions are used in computer science for pattern matching within text (Friedl, 2006), providing a concise and flexible means to find specific sequences of characters within a string with a search pattern. Originally developed for use in theoretical computer science, RegEx is now widely used, from simple string matching to complex text processing and data extraction, using a series of special symbols to define patterns, which can include specific words, numbers, or more complex text structures.

The formulaic (monosaccharides/linkages as building blocks) yet nonlinear nature of glycans requires an adaptation of RegEx for optimal usage. Our RegEx framework closely mimics that developed for text and supports ambiguity, modifiers, quantifiers, greedy and lazy execution, as well as lookahead and lookbehind. A summary is shown in Fig. 1A, with further explanations within glycowork (https://bojarlab.github.io/glycowork/motif.html#regex), since our RegEx system is contained within our open-source Python package glycowork (Thomès *et al*., 2021) (v1.1+) as the new *glycowork*.*motif*.*regex* module, ensuring long-term maintenance. This allows easy access to probe glycan sequences with RegEx patterns via the *get_match* function. RegEx also enhances existing glycowork functionalities, such as specifying RegEx patterns within motif highlighting capabilities of GlycoDraw (Lundstrøm, Urban, Thomès, *et al*., 2023) (Fig. 1B) or adding patterns to motif annotation and network highlighting functions.

**Figure 1.**
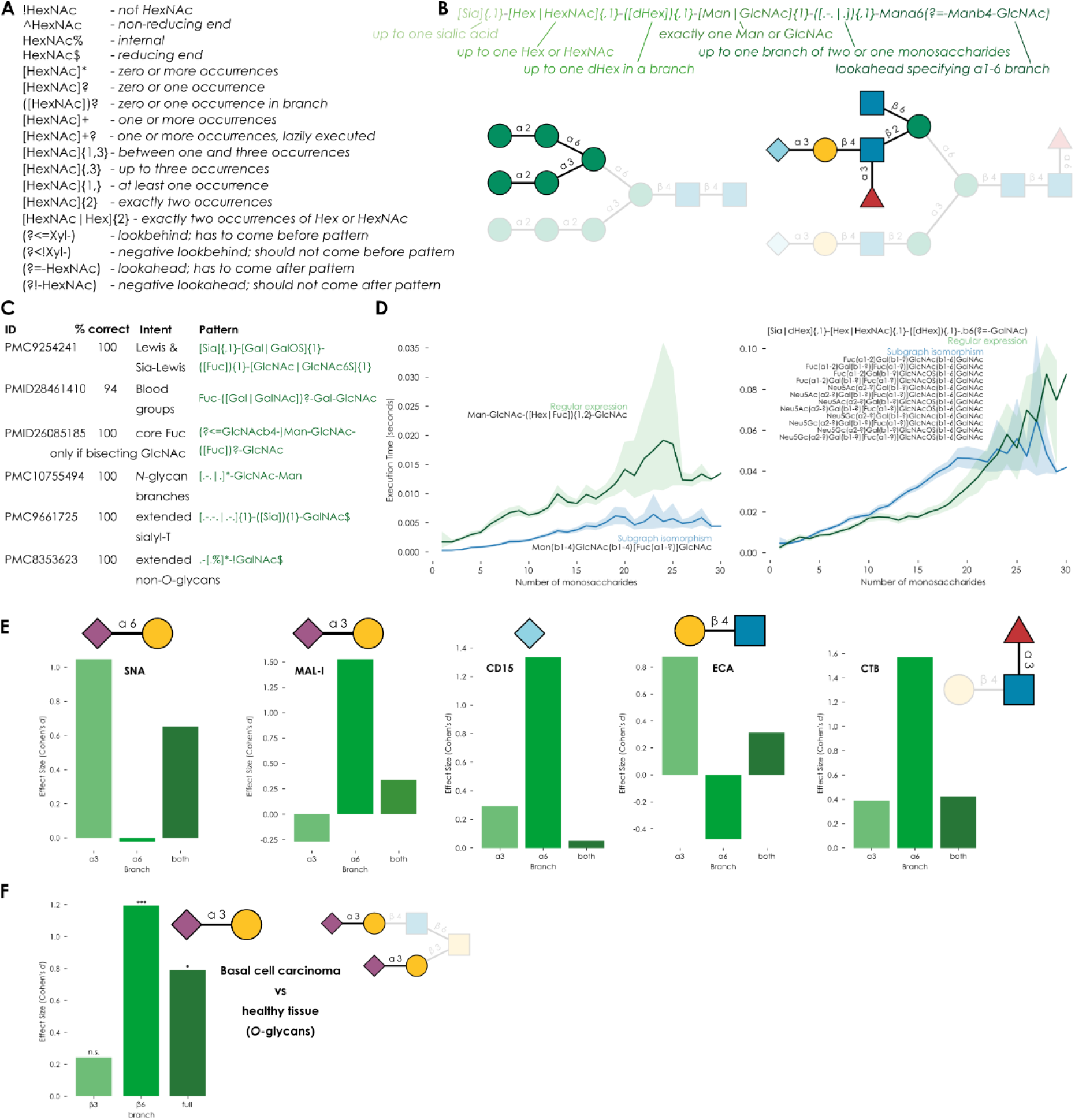
Developing a regular expression system for glycans. **A)** Modifiers and quantifiers for the glycan RegEx system. **B)** An example query designed to extract the α1-6 branch of *N*-glycans, illustrated with GlycoDraw via the “highlight_motif” functionality. **C)** Average accuracy of glycan RegEx queries across multiple datasets, checked manually. **D)** Speed comparison of glycan RegEx query versus normal motif search for core fucosylation or β1-6 branch in *O*-glycans, against all human glycans up to 30 monosaccharides within glycowork. **E)** Sequence contexts of the α1-3 branch, α1-6 branch, and the entire glycan were extracted from the asymmetric *N*-glycan array data (Li *et al*., 2020) and were analyzed with the *get_pvals_motifs* function of glycowork to establish branch- dependent enrichment patterns for various lectins. **F)** Differential glycomics expression of mono- and disaccharide motifs from the β1-3 and β1-6 branch of basal cell carcinoma (Möginger *et al*., 2018) were analyzed with the *get_differential_expression* function of glycowork to identify how branch-specific motifs drive overall patterns.

Our RegEx system first chunks the provided pattern into homogeneous modules (Fig. 1B). Segments without modifiers or quantifiers (but supporting structural ambiguity) are immediately used for subgraph isomorphism operations to detect their location(s) within the glycan graph as prepared by glycowork. Complete wildcards can be specified with “Monosaccharide” or “.”, and linkages can be specified if desired (e.g., “Mana6” or “Galb3/4”) Segments with modifiers or quantifiers are processed into dictionaries of type (glycan substructure: allowed counts), followed by subgraph isomorphisms of the substructures. Glycan branches are indicated by parentheses. Lookahead and lookbehind operations include the specified sequence in matching operations but exclude their node indices from extracted matches. All this is followed by an iterative procedure of tracing a path through the matches of the individual segments, such that all requirements of the regular expression are fulfilled (e.g., minimum/maximum counts) and the match represents a connected portion of the glycan graph. Our RegEx system by default aims for a greedy match, akin to normal RegEx, yet lazy execution is supported by a “?” modifier (e.g., “+?” instead of “+”).

We designed our glycan RegEx system to be functionally analogous to standard RegEx formulation in Python, though we hasten to add that standard RegEx could not easily be used to interrogate glycans in a similar manner, as it would only operate on the string level and fall prey to ambiguities of glycan string nomenclatures, which we avoid with graph operations. An example query (Fig. 1B) illustrates the potential of the herein developed method. Applied to a variety of real-world datasets, followed by manual inspection of correct extraction, yields an average accuracy of 99% (Fig. 1C), for the analyzed datasets.

Simpler glycan RegEx operations are computationally more expensive than motif searches within glycowork (Fig. 1D), yet more complex queries, requiring many conventional motif searches, gain in computational efficiency in comparison. Glycan RegEx allows us to extract larger sequence contexts and use them in downstream analyses, such as the investigation whether sequence patterns of the α1-3/α1-6 branch in *N*-glycans differ across conditions. This presents a non-trivial endeavor as the structural heterogeneity of *N*-glycans does not currently enable a generalizable and easy way to extract α1-6 branches across many sequences. We applied our RegEx strategy to glycan array data from asymmetric glycans, to probe branch-specific binding of well-known lectins (Fig. 1E). This confirmed strong preferences of SNA for Neu5Acα2-6 on the α1-3 branch and MAL-I recognizing Neu5Acα2-3 on the α1-6 branch. We further find enhancement of ECA binding by LacNAc repeats only on the α1-3 branch, and strong preferences for Neu5Gc binding of the αCD15 antibody, only on the α1-6 branch.

To illustrate that this extends beyond glycan array analysis, we engaged in differential glycomics expression analysis (Lundstrøm, Urban, and Bojar, 2023) to find out whether features of specific *O*-glycan branches are differentially expressed in a skin cancer dataset (Möginger *et al*., 2018) (Fig. 1F). We then contrasted this with the current approach of analyzing entire glycan sequences for differential expression and conclude that the increase in α2-3 linked sialylation in cancer, seen in the overall dataset, is entirely driven by the β1-6 branch in this dataset, and hence exclusive to core 2/4/6 structures.

In summary, we envision that a glycan-specific RegEx system will facilitate analyzing glycans and their changes in disease. As this method allows for the analysis of unprecedented and arguably arbitrary motif complexity, we expect the discovery of new patterns in glycan dysregulation. We are also convinced that this approach further strengthens the usefulness of the glycowork ecosystem and will, in general, empower glycoinformatics workflows.

## Author contributions

Conceptualization: D.B., Funding Acquisition: D.B., Resources: D.B., Software: A.R.B., D.B., Supervision: D.B., Visualization: A.R.B., D.B., Writing—Original Draft Preparation: D.B., Writing—Review & Editing: A.R.B., D.B.

## Data availability

Code and documentation are available via glycowork and https://github.com/BojarLab/glycowork/blob/master/glycowork/motif/regex.py.

